# The Biomechanics of the Locust Ovipositor Valves: a Unique Digging Apparatus

**DOI:** 10.1101/2021.12.22.473831

**Authors:** Rakesh Das, Shmuel Gershon, Benny Bar-On, Maryam Tadayon, Amir Ayali, Bat-El Pinchasik

## Abstract

The female locust has a unique mechanism for digging in order to deposit its eggs deep in the ground. It utilizes two pairs of sclerotized valves to displace the granular matter, while extending its abdomen as it propagates underground. This ensures optimal conditions for the eggs to incubate, and provides them with protection from predators. Here, two major axes of operation of the digging valves are identified, one in parallel to the propagation direction of the ovipositor, and one perpendicular to it. The direction-dependent biomechanics of the locust major, dorsal digging valves are quantified and analyzed, under forces in the physiological range and beyond, considering hydration level, as well as the females’ age, or sexual maturation state. Our findings reveal that the responses of the valves to compression forces in the specific directions change upon sexual maturation to follow their function, and depend on environmental conditions. Namely, in the physiological force range, the valves are resistant to mechanical failure. In addition, mature females, which lay eggs, have stiffer valves, up to roughly nineteen times the stiffness of the pre-mature locusts. The valves are stiffer in the major working direction, corresponding to soil shuffling and compression, compared to the direction of propagation. Hydration of the valves reduces their stiffness but increases their resilience against failure. These findings provide mechanical and materials guidelines for the design of novel non-drilling excavating tools, including 3D-printed anisotropic materials based on composites.

**Statement of significance:** The female locust lay its eggs underground in order to protect them from predators and to provide them with optimal conditions for hatching. In order to dig into the ground, it uses two pairs of valves: The ventral pair is plugged as a wedge, while the dorsal pair performs the digging of the oviposition tunnel. We study the mechanical response of the digging valves, depending on age, hydration level and direction of operation. Our findings show that during the course of roughly two weeks in the life of the adult female, the digging valves become up to nineteen-fold stiffer against failure, in order to fulfill their function as diggers. While hydration reduces the stiffness, it also increases the resilience against failure and renders the valves unbreakable within the estimated physiological force range and beyond. The digging valves are consistently stiffer in the digging direction than in the perpendicular direction, implying on their form-follows-function design.

**Graphical Abstract:** 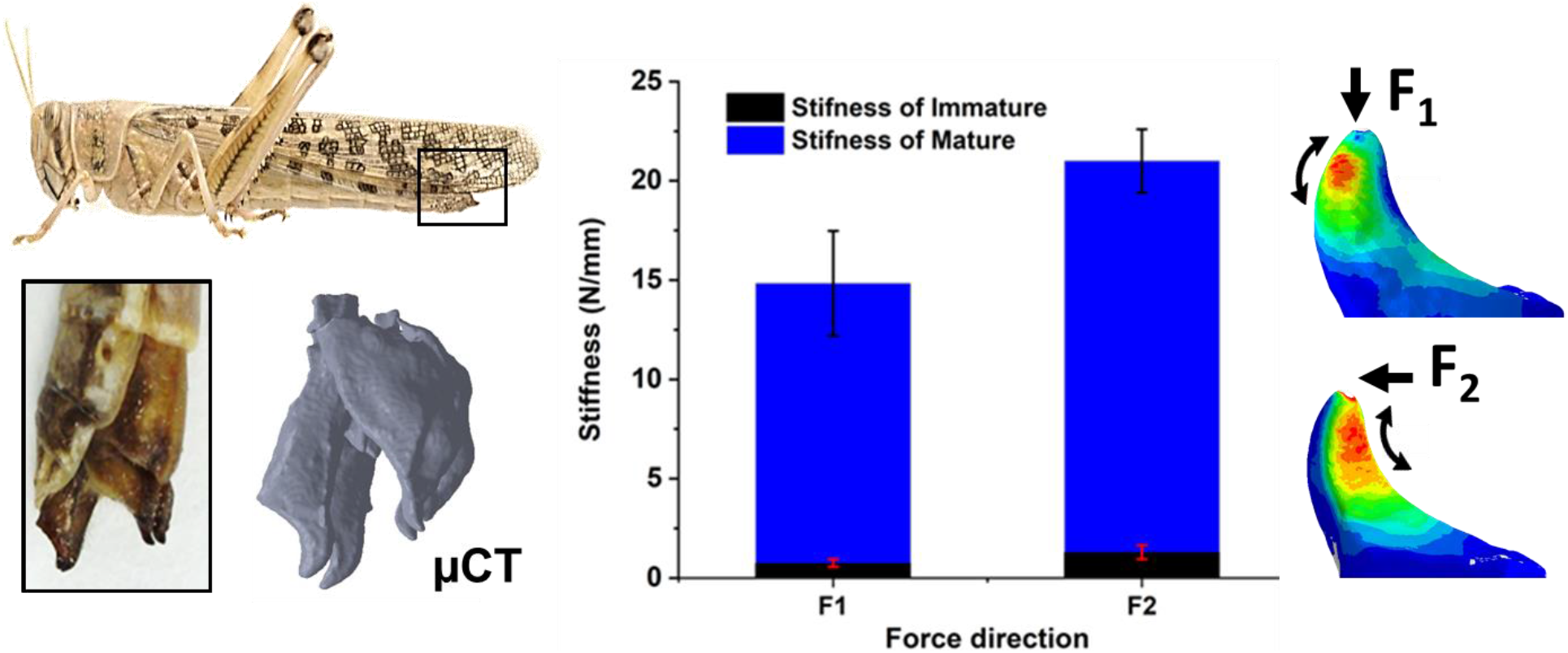

## 1. Introduction

Insects have evolved excellent biological features that promote subsurface exploration and even motion in the subterranean environment^1–4^. The relations between structure, mechanical properties, and function are manifested in various successful digging mechanisms in nature^5–7^. Some of them rely on enlarged front limbs^7^, while some rely on specialized body movements^4^. Several studies presented bioinspired diggers based on such digging apparatuses^8–11^. One promising example is found in the methods and structures used in insect, specifically grasshoppers, oviposition or egg-laying^12–16^. These methods rely on ad-hoc digging apparatuses. Egg laying is a central aspect of the reproductive biology of insects. The deposition of eggs in a carefully selected spot, in or on a carefully selected substrate, represents a major decision and a crucial act that the female insect executes to ensure the survival of its progeny. Evolution and natural selection have, therefore, acted to perfect the related mechanisms and maximize the chances of successful oviposition. One manifestation of the above is a striking diversification of the dedicated apparatuses in different insects, ensuring the extreme and intricate adaptations of the related body structures to their function and to the selected environment and substrate^13^.

The ovipositor of grasshoppers and locusts is a highly specialized structure consisting of two pairs of shovel-shaped cuticular valves, a ventral and a dorsal pair, extending beyond the distal end of the female abdomen (**Fig. 1a**). The valves are hinged at their bases to each other and to a prominent pair of internal apodemes, a ridge-like ingrowth of the exoskeleton, serving the large supporting muscles^17^. These structures are used to dig a deep and narrow chamber in the ground for egg burial, to manipulate the eggs, and to assist in capping the egg-pod with froth (Fig. 1b). During oviposition, the ovipositor valves undergo rhythmic cyclical opening and closing, retraction and protraction movements^16^. These movements are produced by the contractions of ten pairs of muscles innervated by the terminal abdominal ganglion. Grasshoppers and locusts are rare among insects in having ovipositor valves that work by opening and closing movements rather than by valves sliding upon each other^15^. The locust ovipositor valves exert different forces needed for digging, clearing debris from the digging path, and for the hyperextension of the female’s abdomen during the oviposition. These tasks are divided between the valves: the ventral ones are mostly responsible for the pulling action, while the dorsal valves are used for clearing debris.

**Fig. 1.**
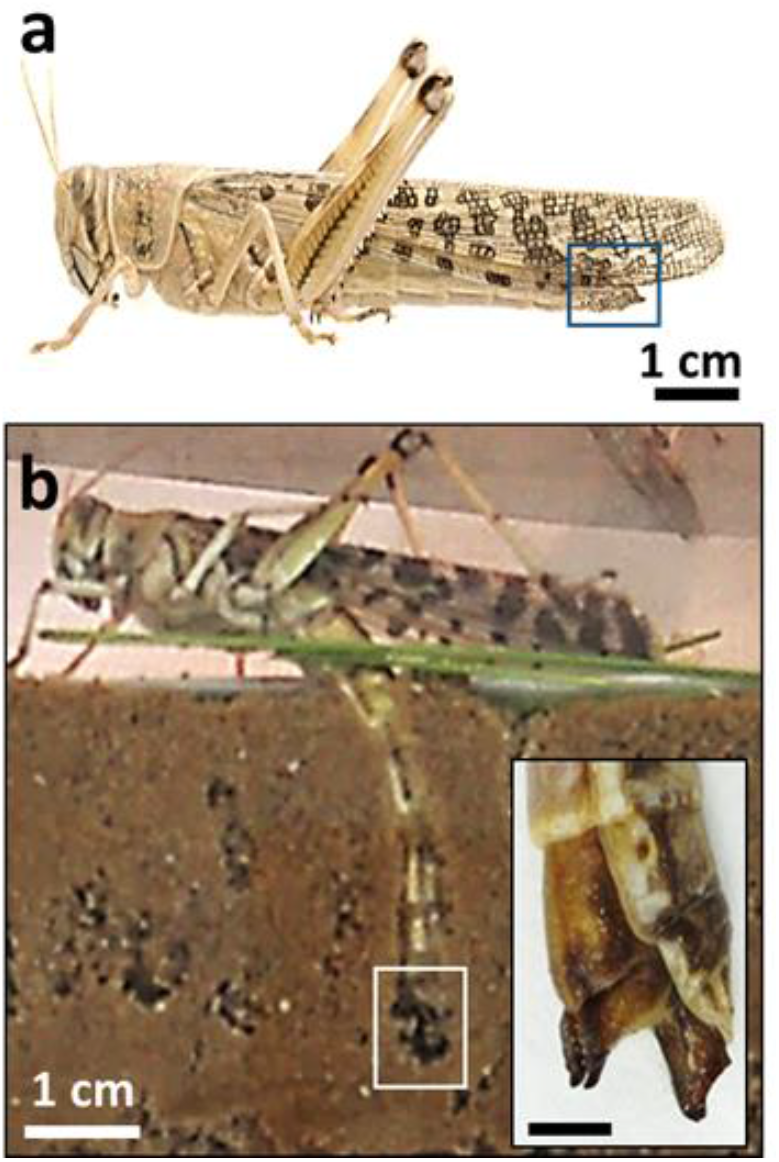
The female locust oviposition underground. a) The ovipositor of the female locust at the tip of the abdomen (noted by the blue square). b) A locust female during oviposition, extending its abdomen into the ground while digging. Inset: higher magnification of the two pairs of digging valves in their open state, scale bars correspond to 5 mm.

In newly hatched locust females, the valves are visible as paired outgrowths of the posterior margins of the last abdominal segments. They enlarge and differentiate in a series of steps across the five larval stages, and by the third larval instar, the valves adopt the adult ventral/dorsal orientation. In the fourth and fifth larval stages, the valves continue to enlarge with each molt, such that they extend beyond the abdomen tip at the final molt to adulthood.

The soft valves enlarge further and become densely sclerotized during two or three more weeks while the animal reaches sexual maturity^18–20^. How the mechanical properties of the locust ovipositor valves change with the adult age, and specifically during sexual maturation to facilitate the digging for oviposition, remains an open question.

In order to understand how the shape and mechanical response of organs serve their function in nature, specifically in arthopods^21–24^, and to develop design and mechanical guidelines for bioinspired diggers^25^, the quantification of biomechanical factors is essential^26–28^. Namely, the physiological force range, force-deformation behavior, maximal force before failure, and the dependence of the mechanical response on direction. In addition, in the case of insect cuticles, one of the major factors influencing the mechanical properties is the hydration state^29–31^.

In this study, we quantify the mechanical response of the locust ovipositor dorsal valves, depending on age (sexual maturation state), hydration, and force direction. We focus on the dorsal valves since their shovel-shaped apex region plays the primary role in shuffling and compressing the soil. We explore two major force directions, corresponding to the direction of propagation and direction of excavation, i.e., to the protraction and opening of the valves, respectively. We approximate the physiological force range in which the valves operate and discuss the mechanical stability of the valves within this range and beyond. We use compression mechanical tests to reveal the relations between an applied force and deformations of the valves, depending on hydration level. In addition, we quantify the maximal forces the valves can withstand in the two above-mentioned directions. We quantify how the structural stiffness and maximal load-bearing force is altered with age (i.e. sexual maturation), and adapt to the mechanical needs during oviposition. Finally, we use Finite Element (FE) simulations, based on 3D micro-computed tomography scans, in order to model the load distribution in the valves during excavation, and we elucidate the failure mechanism of the valves in experiments. Our findings shed light on the mechanical requirements for fulfilling the function of digging underground and can inspire the development of synthetic 3D-printed valves with direction dependent improved mechanics.^32–34^

## 2. Materials and methods

### 2.1. Sample preparation

Locusts were obtained from our desert locust, *Schistocerca gregaria*, colony at the School of Zoology, Tel Aviv University. Ovipositor valves were dissected from females of two different groups: 1) sexually mature females (age ≥ 30 days post molt to adult, with oviposition history) and 2) younger, pre-mature females (age = 7-9 days post molt). After extraction of the valves, they were cleaned and kept in a sealed tube at room temperature (25 °C). If required, the valves were preserved in a freezer (−20 °C) for a maximum of 48 h. To avoid desiccation, the samples were surrounded by wet cotton and sealed with Parafilm. These preparation and preservation procedures were previously demonstrated to have no significant influences on the samples’ biomechanical properties^35^. To dehydrate the samples, they were sequentially immersed in ethanol with increasing concentrations (30 %, 50 %, 75 %, and 100 %).

### 2.2. Micro-computed tomography (μCT) scanning based 3D-model

The ovipositor valves were scanned by a μ-CT machine (Easytom, RX-Solutions, Chavanod, France) at an isometric voxel size of 5 μm at 125 μA and tube voltage of 80 kV, equipped with a micro-focus tube (XRay150, RX-Solutions) and a flat panel detector (CsI scintillator). The 3D model was built after the reconstruction and segmentation of 3D scanned images based on grey value using Simpleware Scan IP software (Synopsys, CA, USA). In this segmentation process, different built-in tools (median filter, island removal, and flood fill) were utilized to reduce the noise. The segmented 3D model meshed in an FE model in the advanced meshing module of Simpleware Scan IP with tetrahedral elements (C3D4). The number of elements was 74802, in the model of the dorsal valve tip. This meshed 3D model was further used for numerical simulation. A Mesh convergence study was performed to make sure that the element size has no influence on the analysis.

### 2.3 Bending cantilever experiments

The experimental setup (**Fig. 2a**) included a sample holder, in which the locust abdomen is fixed while the valves are free to perform the digging motions. A steel cantilever was located as to barely touch the dorsal valves. We took advantage of the well-established fact that the oviposition motor program can be activated by releasing the local control system from descending inhibition, i.e. by transection of the ventral nerve commissures caudal to the abdominal ganglia^12^. The digging movements of the valves induced displacement of the beam and this displacement was imaged using a camera (Panasonic DC-S1 with Sigma 70 mm *f*/2.8 DG Macro lens). The generated force is considered a point load, as the diameter of the dorsal tip is considerably smaller than the deflecting edge of the beam. ImageJ freeware was used to analyze the images and extract cantilever deflection (*δ*). The force was extracted from the cantilever deflection via *p* = 3*δEI/L*^337^, where *L* = 5.2 *cm* is the cantilever length, *I* = *WT*^3^/12 is the cantilever moment of inertia with *W* = 0.27*cm* and *T* = 0.1 *cm*, and *E* = 180 *GPa* is the Young’s modulus of the cantilever’s material..

**Fig. 2.**
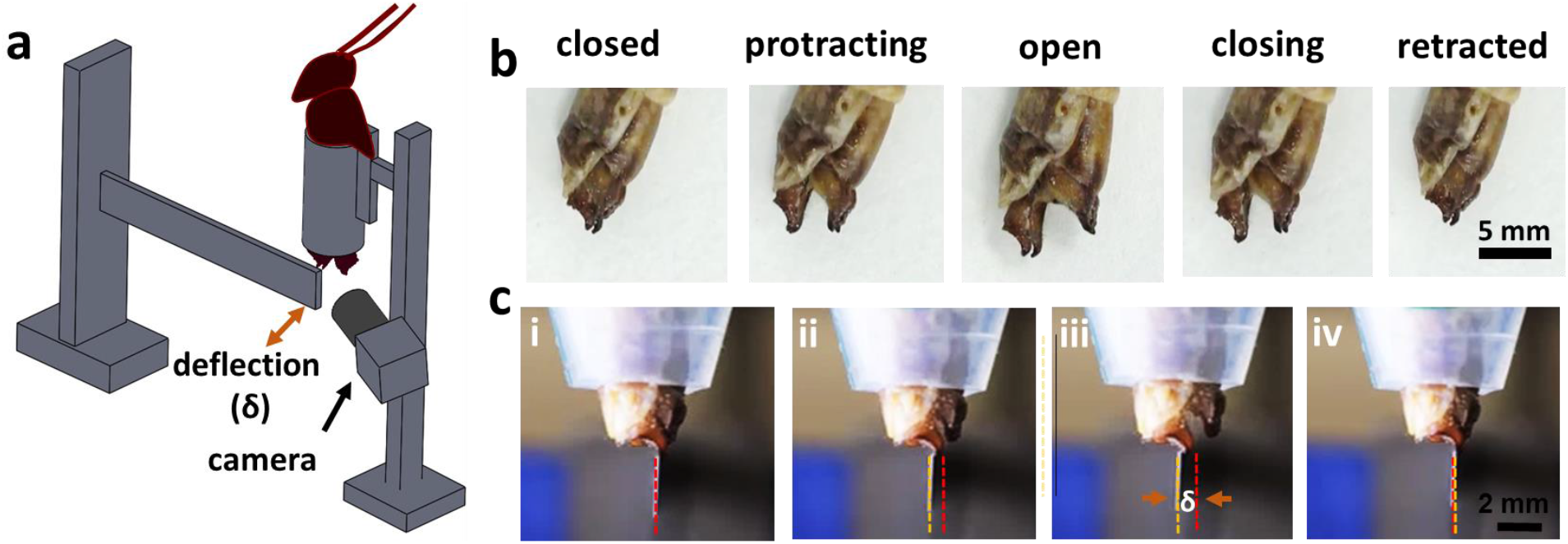
Operation and estimated physiological force range of the locust valves. a) An illustration of the experimental bending beam setup, used to measure the approximated physiological force range exerted on the dorsal valves during digging. b) The different states of valves during digging. c) The cantilever bending experiment. The red dashed and the dashed yellow lines correspond to the original and deflected positions of the beam, respectively. *δ* is the cantilever deflection.

### 2.4 Tip-loading mechanical tests

The shafts of the valves were embedded in epoxy glue onto a screw-head, exposing only the apex region. The screw was mounted on a customized grip of a mechanical testing system (TA Instruments, USA) in order to make the base of the samples immobile when the tip of the samples touched the compression plate. A preload of 0.2 N was applied through the compression plate on the tip. A constant displacement rate of 0.005 mm·s^-1^ was applied on the tip, with a total displacement of 0.7 mm. The corresponding force was recorded by the load cell of the system. The two reference directions of the applied compressive force were chosen based on the functioning of the valves during the excavation. The force-displacement curves (*F* – *d*) were obtained from this experiment, in which the initial slope represents the elastic stiffness, and the compressive breaking force was calculated from the maximum of these curves.

### 2.5 Scanning Electron Microscopy (SEM)

The fractured specimens of the ovipositor valves were imaged using a Scanning Electron Microscope (GeminiSEM300, Zeiss, Oberkochen, Germany). The fractured specimens were obtained after compression mechanical test (see above) in two reference force directions. Before scanning, the samples were sputter-coated using Polaron sputter coater, model SC7640, Au-Pd target (Polaron England).

### 2.6 Finite Element Simulation for the mechanical behavior of the valves

The 3D meshed models of the dorsal and ventral valves pair were exported to Abaqus-FEA package (version 6.14; ABAQUS Inc., CA, USA) for analysis through finite element simulation. The average material properties determined from nanoindentation (see supporting information, *Nanoindentation tests*, **Fig. S2**) were applied in these models. During the simulation, the valves are considered linear elastic and isotropic. The boundary conditions for the compression mechanical test were applied. The base of the valves were fixed (all translational and rotational degrees of freedom were constrained) and the force determined from the compression mechanical experiment was applied at the tip of the valves. The resulting principal stress distributions were analyzed.

## 3. Results

### 3.1 Physiological force measures of the valve

To quantify the physiological force range that the female locust’s digging apparatus operates in, we performed a bending cantilever experiment (**Fig. 2**).

We took advantage of the well-established fact that the oviposition motor program can be activated by releasing the local control system from descending inhibition, i.e. by transection of the ventral nerve commissures caudal to the abdominal ganglia^12^. This results in induction of the ovipositor digging movements (or fictive digging movements) out of the context of egg laying, and outside of the substrate.

Fig. 2a illustrates the experimental setup, in which the locust abdomen is fixed while the valves are induced to perform the digging motions. As noted above, during digging, the valves go through cyclic opening and closing (Fig. 2b). Initially, the valves are in their closed/retracted state. The ovipositor then protracts, opens and closes, and retracts in a complete cycle^37^. A full cycle, under the conditions used in this experiment, corresponds to approximately 3-4 seconds. This is in agreement with the time span of a full digging cycle underground (see supplementary **video V1**).

A representative image sequence of the bending beam experiment is shown in Fig. 2c (see supplementary **video V2**). Upon the opening movements of the valves, the cantilever experiences deflection (Fig. 2c ii,iii), and the force is calculated (see Materials and Methods). Any abdominal movements beyond those of the valves are restricted to cancel potential influence on the beam deflection. This calculation yielded a force corresponding to 0.85 ± 0.15 N (n=10) in the direction of opening (perpendicular to the propagation). Since the experiment is conducted in air, the actual forces exerted by the intact female locust may differ during real operation underground (due to resistance or feedback from the granular matter). Nevertheless, this measurement and calculation provides an estimate of the physiological forces for later discussion and corresponds to the maximal excavation force applied by the female.

### 3.2 Mechanical response of the locust valves to tip-loadings

We quantified the mechanical response of the locust dorsal valves to normal and lateral tip-loadings, which correspond to its propagation and digging actions, respectively (**Fig. 3**).

**Fig. 3.**
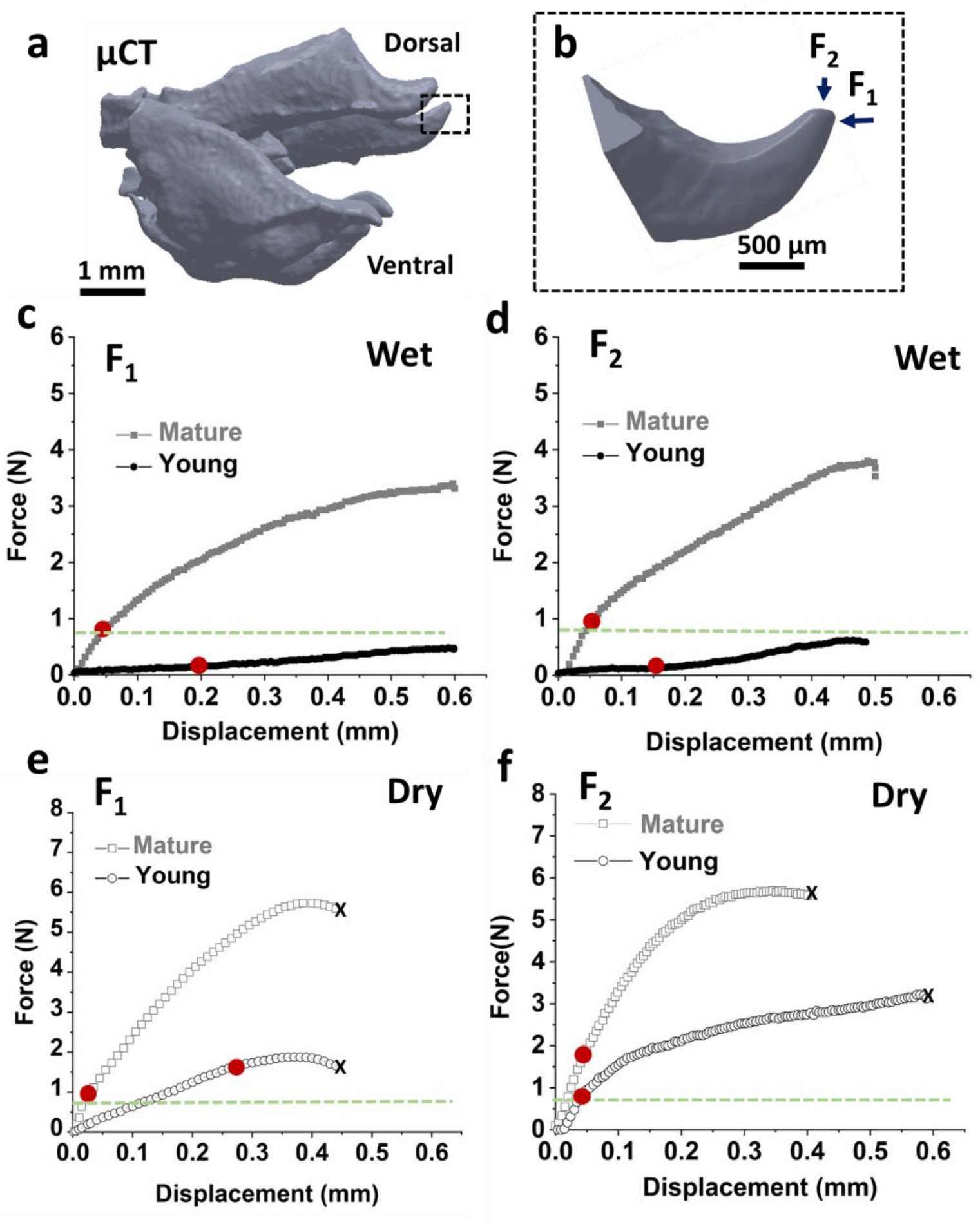
Mechanical response of the locust valves under uniaxial compression. a) μCT reconstruction of the entire dorsal and ventral valves, and b) a higher magnification of the dorsal valve’s tip. Two major force axes are noted: in the direction of propagation, F_1_, and direction of excavation, F_2_. c-d) Representative force-displacement curves of mature (grey) and younger pre-mature (black) hydrated (wet) valves under forces F_1_ and F_2_. Representative force-displacement curves of mature (grey circles) and young (black circles) dehydrated (dry) valves under forces F_1_ and F_2_. The end of the linear range, which represents the elastic stiffness response, is marked with red dots on each of the curves in the hydrated and dehydrated conditions, respectively. The x indicates the point-of-failure (occurs only in dry state). The green dashed lines represent the maximum of the physiological force range.

Fig. 3a shows a μCT scan reconstruction of the entire dorsal and ventral valves. Fig. 3b shows a higher magnification of the dorsal valve’s tip under load in the two major axes: the direction of propagation, F_1_, and the direction of excavation, F_2_.

Representative force-displacement curves are shown in Fig. 3 c-f. We quantify the response both in the estimated physiological range of up to roughly 0.85 N, marked by green dashed lines in the plots, and further until failure of the valves. We compare valves of mature (grey) and younger pre-mature (black) locusts and quantify the role of hydration in their mechanical stability in the two force directions. Four main points are evident: i) valves of mature locusts are significantly stiffer than those of young locusts, ii) valves are stiffer in the *F*_2_ direction, in comparison to *F*_1_, iii) hydrated (wet) valves are less stiff but more resilient to failure, both in the physiological operation range and beyond, and iv) hydration influences mostly the mechanical properties, namely stiffness, of the younger locusts in comparison to the mature ones.

**Fig. 4** summarizes the mechanical response of the dorsal valve’s tip depending on age and hydration level (Fig. 4a,b) and presents the breaking force of dehydrated valves in both directions (Fig. 4c).

**Fig. 4.**
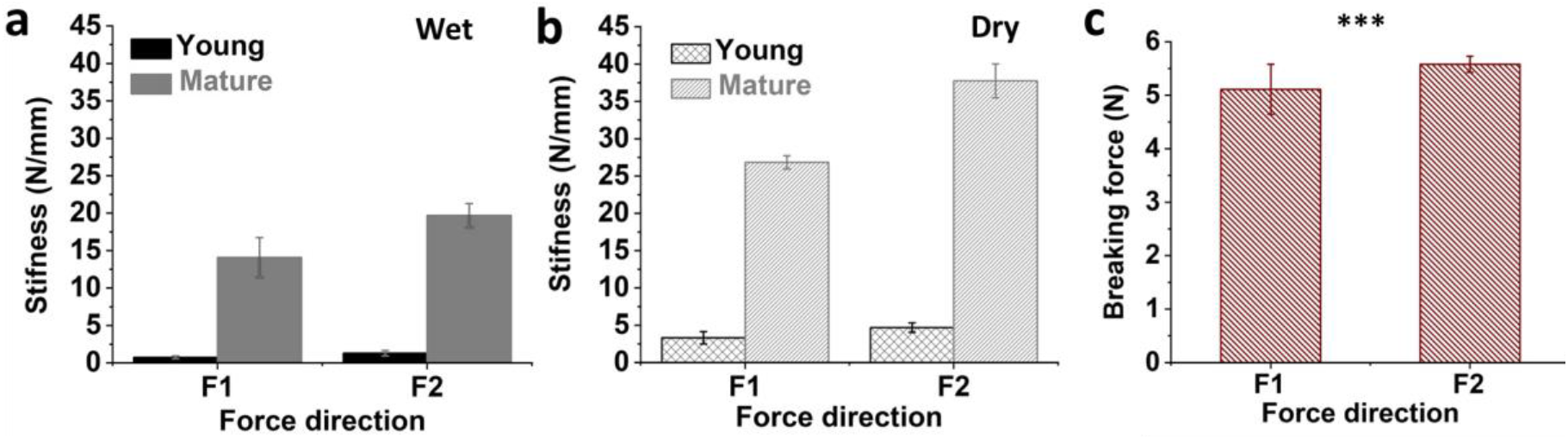
The age, hydration, and direction-dependent mechanical response of the locust valves. The structural stiffness of the dorsal valve’s tip of young and adult locusts in a) hydrated (wet) and b) dehydrated (dry) conditions. c) Breaking force of the valves in dehydrated conditions. (***: P ≤ 0.01, Student’s t-test).

In the physiological range, the structural stiffness of the dorsal valves of a mature locust, in the wet state in F_1_ direction, corresponds to 14.08 ± 2.64 N·mm^-1^ (n=10), almost nineteen-fold the stiffness of the valves of the pre-mature locusts, with 0.45 ± 0.12 N·mm^-1^ (n=10) (Fig. 4a). The stiffness was calculated from the initial linear slope of the force-displacement curve. Interestingly, wet valves did not break under compression, even for displacements of 0.6 mm, corresponding to forces of up to approximately 4 N in the mature locusts and 0.6 N in the young locusts. However, an irreversible residual deformation was visible in all hydrated samples (**Fig. S1** in the Supporting Information). In force direction F_2_, the stiffness of the wet mature locust valves in the physiological range corresponds to 26.83 ± 0.89 N·mm^-1^ (n=10), almost double the stiffness in F_1_ direction of 14.08 ± 2.64 N·mm^-1^ (n=10). In addition, in the F_2_ direction, the wet valves of the mature locusts are significantly stiffer than the valves of the wet young ones, with 19.69 ± 1.60 N·mm^-1^ and 1.30 ± 0.35 N·mm^-1^ for the mature and young locusts, respectively (Fig. 4a).

While the hydration level of the valves is unknown in the living locust, we observe a strong dependence of the mechanical properties on the hydration state. The stiffness is, as expected, higher for dry valves, corresponding to roughly twice the stiffness of the wet ones for both force directions in mature locusts (Fig. 4a,b). Namely, 26.83 ± 0.89 N·mm^-1^ in comparison to 14.08 ± 2.64 N·mm^-1^ in the F_1_ direction for mature locusts, for dry and wet valves, respectively, and stiffnesses of 37.75 ± 2.27 N·mm^-1^ in comparison to 19.69 ± 1.60 N·mm^-1^, in the F_2_ direction, for dry and wet conditions, respectively. Also in the dehydrated valves, the structural stiffness in the F_2_ direction is higher than in the F_1_ direction, with 19.69 ± 1.60 N·mm^-1^ (n=10) in comparison to 14.08 ± 2.64 N·mm^-1^, respectively.

It is well established that insects’ cuticles lose water during sclerotization^38,39^. With our results, we are able to quantify the influence of hydration between young and mature locusts. For example, for a young locust, the structural stiffness in the F_1_ direction increases from 0.45 ± 0.12 N/mm (wet) to 3.32 ± 0.83 N·mm^-1^ (dry). This corresponds to an increase of roughly 640 % in stiffness. However, for a mature locust, the increase in stiffness is only 90 %, from 14.08 ± 2.64 N·mm^-1^ to 26.83 ± 0.89 N·mm^-1^. Most likely, due to lower water content in the more sclerotized areas of the organ^20^.

While in the estimated physiological force range we did not observe failure of the valves, in either the young or the mature valves, in wet or dry states, we observe the failure of the valves in higher compression forces, within the range of 0.6 mm, corresponding to the maximal displacement in the experiment. None of the valves failed in the wet conditions. However, the valves of the mature locust failed when they were dehydrated under compression forces of 5.58 ± 0.15 N, in the F_2_ direction, in comparison to 5.11 ± 0.47 N, in the F_1_ direction. While these values are far from the physiological conditions, they still maintain the trend of increased mechanical stability in the direction of digging (F_2_).

### 3.3 Mechanical failure

We now focus on the structural position and the mechanism of the mechanical failure of the dorsal valves in their dry state, under forces in the propagation (F_1_) and digging (F_2_) directions (**Fig. 5**). Fig. 5a-d and Fig. 5e-h show optical micrographs of the valves in the initial state and after failure, scanning electron micrographs of broken valves and finite element simulations of the maximal principal stress distribution in the dorsal valves in F_1_ and F_2_ directions, respectively. The fracture topography implies on a brittle material^40^, agreeing with the force-displacement curves of the dry valves (Fig. 3 e,f). Therefore, the criterion for failure corresponds to the maximal tension developing in the valves under external forces^41^.

**Fig. 5.**
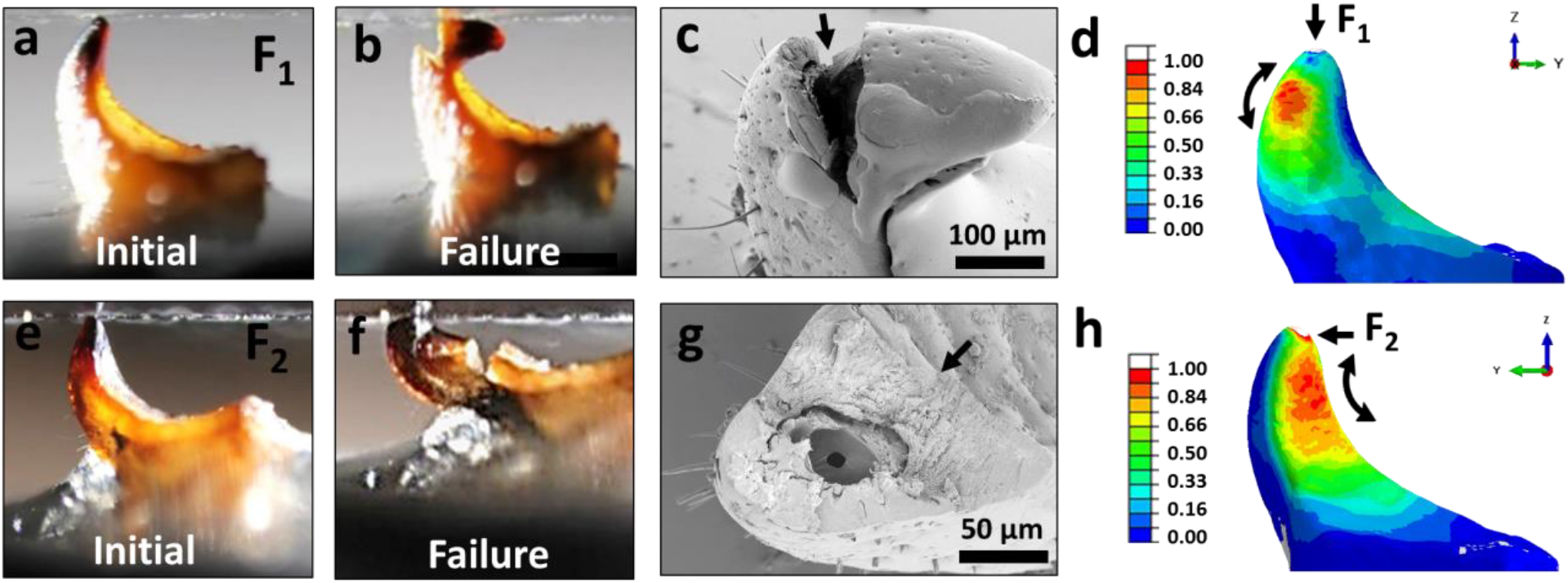
Analysis of the mechanical failure of the locust valves: experiments and simulations. Optical micrographs of a) the valve in the initial state and b) after failure under force in direction F1. c) Scanning electron micrographs of the valves after their mechanical failure, the black arrow indicates the direction of the fracture. d) Simulated normalized maximal principal stress in the valves under force F_1_. Equivalent images and simulations for force direction F_2_ are depicted in e-h.

In order to understand the failure mechanism of the valves, the stresses developing under external loads were modeled. To this end, we mapped the maximal principal stress distribution in the dorsal valves under applied forces in F_1_ and F_2_ directions using Finite Element (FE) simulations. The normalized stresses are shown in Fig. 5d and Fig. 5h, respectively.

For the simulations, we used the μCT scans of the dorsal valves (Fig. 3b). The base of the valves was fixed with full encastre condition, and the forces F_1_ and F_2_ were applied at the tip of the valve with a magnitude of 5 N, corresponding to the breaking force obtained from the compression tests. The forces were applied in the directions corresponding to the compression tests: F_1_ was applied by defining pressure at the valve’s tip in the direction of propagation, and F_2_ was applied by defining surface traction at the tip in the direction of soil removal. The material was assumed to be linear-elastic and isotropic. The elastic modulus was defined as E = 5 GPa, which is the average elastic modulus of mature dry dorsal valves, measured by nanoindentation (see Supporting Information, Fig. S2). Poisson’s ratio was defined to be 0.38.^6^

Fig. 5d presents the convex side of the valves, corresponding to the internal side of the digging system (Fig. 2b), while Fig. 5h presents the concave side, corresponding to the external side in the digging system. The black curved arrows indicate the direction of the tensile principal stress component at the location of its maximal value. During compression under F_1_, the failure of the valve occurs at the convex region (the spine of the valve), as shown in Fig. 5d. We find remarkable agreement between the area of maximal tensile stress in the simulation and the point of failure in the experiment (Fig. 5b,c). We exclude concentrated stresses at load application areas. Similarly, the area of the maximal tensile stress, under F_2_ in the simulation, agrees with the compression experiment (Fig. 5e). In this case, the failure starts from the concave surface, where the maximal tensile stress develops.

## 4. Discussion

The female locust has a unique way of digging in granular material, in which the ground is being compressed to the sides of the burrow rather then removed, and for which the digging valves are remarkably adapted. During underground excavation, the digging valves of the female locust are subjected to forces of approximately 0.85 N in the direction of digging. The stiffness of the valves increases almost nineteen folds until the locust reaches sexual maturation, within two weeks. This remarkable difference shows that the valves’ mechanical stability increases significantly in preparation for their function as digging tools during oviposition.

Considering the force in the physiological operation conditions, the valves are entirely resilient to failure when in their hydrated conditions. Dehydration increases the stiffness of the valves but also renders the valves more prone to fracture under load. Nevertheless, this occurs in forces more than five folds in magnitude than the characteristic forces in the physiological conditions. Regardless of the hydration level, the valves are stiffer in the digging direction in comparison to the propagation direction, implying their design meets the criteria for functioning as resilient digging apparatuses. Dehydrated valves are brittle and fail mechanically due to the development of local tensile forces. Due to the structural gradient of the valves, the failure occurs close to the tip of the valves.

Overall, the valves’ structural gradient, together with the reinforcement of their stiffness upon the locust sexual maturation, and the dependence on hydration, may set clear guidelines for the design of synthetic diggers. First, the structural gradient assures the failure happens closer to the tip, which may allow the organ to operate even if fractured. Second, the stiffness of the valves should be adjusted to the designated granular matter and environmental conditions. Third, water, which functions as a plasticizer, increases the resilience of the digger against fracture and failure. It is, therefore, an interplay between the stiffness and compliance of the material, in order to offer both functionality and resilience.

For mimicking cuticular materials, and specifically digging organs of insects, 3D-printing using composite materials is an attractive direction. Specifically, it provides an ability to alter the mechanical properties of the printed material in specific regions and in specific directions. Considering such 3D-printed composites are very often brittle^42^, we provide simulations of the maximal tension developed in the organ during loads in both the propagation and digging directions. These provide guidelines for the design and possible material enforcement where the maximal tension develops in order to improve the resilience of bioinspired systems against failure^43^.

## 5. Conclusions

The mechanical response of the locust valves corresponds to their biological function and changes dramatically throughout the female locust life cycle. The stiffness of the valves of mature locusts are almost twenty folds higher than that of younger locusts, and the valves’ stiffness is higher in the direction of digging in comparison to the direction of propagation. The hydration of the valves makes them more resilient against failure, showing no breakage in the physiological force range of digging and beyond. The stiffness of the valves of the sexually mature locusts, however, is less affected by dehydration, in comparison to the younger locusts, probably due to lower water content in the mature adult valves. Finally, finite element simulations elucidate why the valves break in specific places, toward the tip, in both force directions. The tensile stress distribution indicates the area in which the tip is most likely to break, as observed in mechanical compression experiments. This is a striking example of the form-follows-function principle in natural structures that can inspire technological innovations and bioinspired mechanical tools for robotics.

## Authors’ contributions

B.E.P., A.A conceived the project idea. A.A. dissected all the locusts and provided the valves. R.D. performed the experiments. S.G. made the simulations. M.T. performed the nanoindentation. B.B.O. contributed in analysis of the simulations and valuable discussions. B.E.P. and A.A. wrote the initial manuscript, and developed figures with input from all authors. All authors provided substantial edits on the final manuscript and gave approval for publication.

## Acknowledgment

We would like to thank Daniel Werner from the Max Planck Institute of Colloids and Interfaces for the μCT data acquisition. We thank Dr. Yoav Lahini from the School of Physics and Astronomy, Faculty of Life Sciences, Tel-Aviv University for assistance in preliminary force measurements. We Thank Prof. Yael Politi from Dresden University, Germany, for valuable comments and fruitful discussions. Rakesh Das acknowledges the funding support of the TATA Trust Post-Doctoral Fellowship-Life Sciences (with Individual Tracks, Ref: 2019-20/PTAU/35).

## Supporting information

**Supplementary video V1:** Digging cycles of the locust in granular matter and in air.

**Supplementary video V2:** Bending beam experiment.

**Fig. S1.**
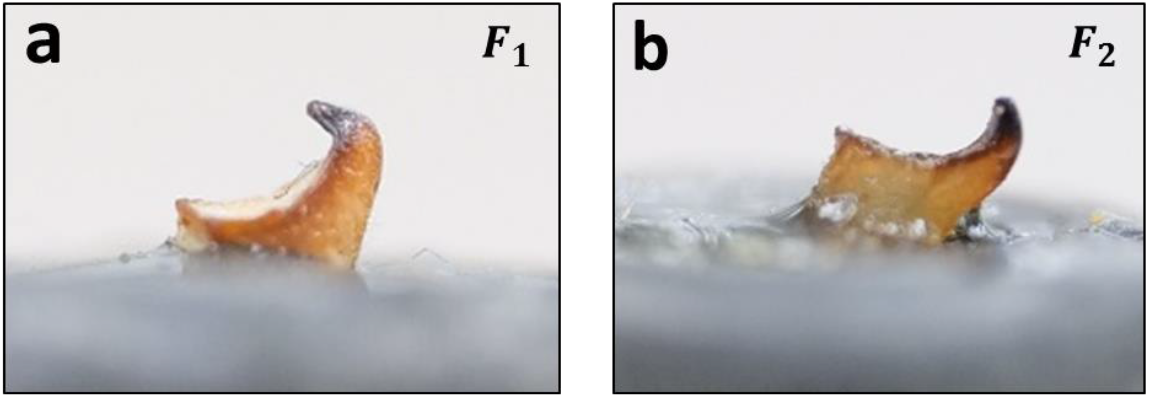
Wet dorsal valves of mature female locust, deformed after compression tests in force directions a) F_1_ and b) F_2_.

### Nanoindentation tests

We used the embedded and finely polished samples to probe the local mechanical properties of the locust valve samples, using a nanomechanical tester, Triboindenter TI-950 (Hysitron, Minneapolis, MN, USA). We used a 30 mN standard transducer with a fluid cell cube cornered diamond tip to quantify the reduced modulus, *E_r_* and hardness, *H*. We set the loading/unloading rate of 60 μN·s^-1^ with a two-second holding time at a peak load of 300 μN. According to the Oliver and Pharr method^1^, load-displacement curves were analyzed towards extraction of reduced modulus, *E_r_*, and hardness, *H*.An array of indents was performed over the overall cross-section of the sample in both dry and hydrated conditions (fully soaked). Two different preloads, 2 and 5 μN, were used to accurately “sense” the sample’s surface in dry and hydrated conditions, respectively. Data analysis was done using OriginPro software (OriginLab Corporation, MA,USA). Values are reported in mean± SD format.

**Fig. S2.**
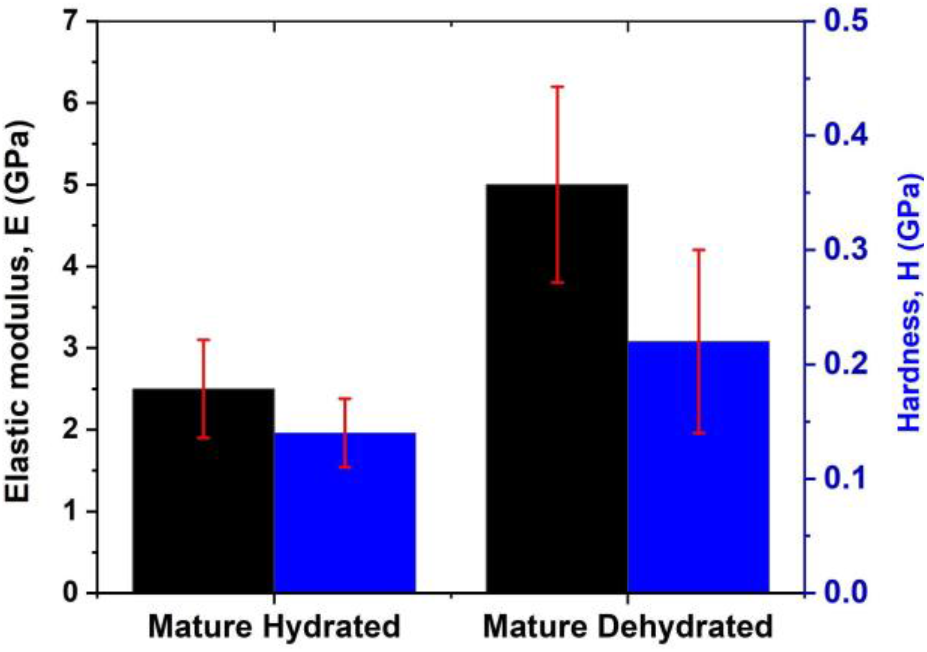
Mechanical Properties of the dorsal valves of a mature female locust, obtained by nanoindentation.

